## Supplementary Information for "Sequence Complexity and Monomer Rigidity Control the Morphologies and Aging Dynamics of Protein Aggregates"


### I. DYNAMICS OF MORPHOLOGICAL TRANSITIONS

We show the morphological transition of aggregates in time at in stiff chains for $H_3 = 1.58$ and $H_3 = 4.75$ in Fig.S1. The corresponding results for $\epsilon_b = 0$ are shown in Fig. 2d in the main text. Higher nematic order parameters for $H_3 = 1.58$ illustrate the fibril formation. In contrast, the $H_3 = 4.75$ sequence at is disordered with low nematic order. At $\epsilon_b = 4$, the value of the nematic order parameter is non-zero (left panel in Fig.S1). The conformational fluctuations are greater in the high-complexity sequence compared to the low-complexity sequence. There is signature of aging for both the high and low complexity sequences for the higher rigidities.

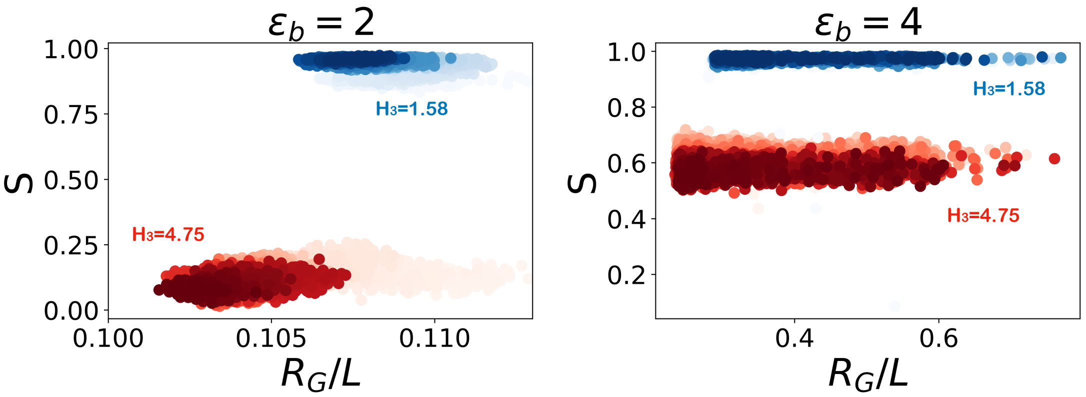


FIG. S1: **Morphological transitions as a function of time for $\epsilon_b = 2$ and $\epsilon_b = 4$:** Trajectories illustrate the evolution of the morphology of aggregates. The $x$-axis represents the radius of gyration of aggregates, $R_G$, normalized by the length of the periodic box, $L = 100\sigma$. The $y$-axis shows the nematic order parameter, $S$. The opacity of the dots indicates progression in time, with denser dots corresponding to more recent times. Blue and red dots correspond to $H_3 = 1.58$ and $H_3 = 4.75$, respectively.

### II. PAIR MEAN SQUARED DISPLACEMENT FOR HIGHER RIGIDITIES

We calculated the pMSD for higher rigidities: $\epsilon_b = 2$ and $\epsilon_b = 4$. In accord with the results in the main text (Fig.5-6), we find evidence for aging dynamics in pMSD for higher rigidities (Fig.S2-S3).

### III. CONNECTION TO THE TRAP MODEL

The theory of aging protein in condensates [S1] is based on the trap model, which first proposed to describe a similar phenomenon in spin glasses [S2, S3]. In essence, aging is attributed to the diminishing fraction of diffusive elements, $P_u(t)$, as the condensate ages, leading to slow dynamics. Although the increased rigidity of monomers adds to the complexity of aging protein condensates, our simulations show that key aspects of the trap model hold. The


*Electronic address:

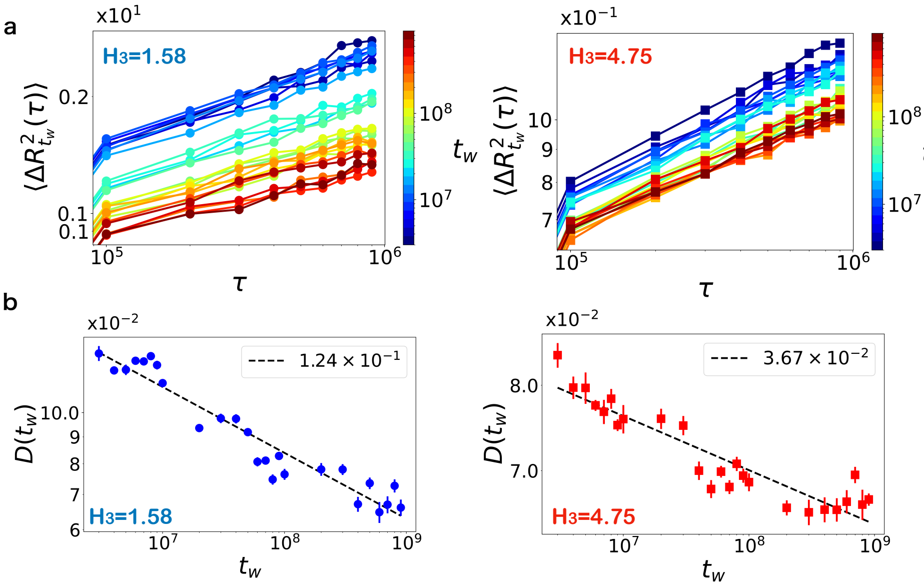


FIG. S2: **pMSD for $\epsilon_b = 2$:** (a) pMSD for low complexity sequence ($H_3 = 1.58$, left) and high complexity sequence ($H_3 = 4.75$, right). The color gradient indicates the system's age ($t_w$). In contrast to Fig.3 in the main text, pMSD for both $H_3$ values depend on the waiting time, $t_w$. (b) The generalized diffusion coefficient $D(t_w)$ as a function of the age $t_w$ obtained by fitting Eq.(5) in the main text to Fig.S2(a). The error bars for $D(t_w)$ are the standard errors. The dashed line is a power-law fit with exponent $\alpha = (1.24 \pm 0.16) \times 10^{-1}$ for $H_3 = 1.58$ and $\alpha = (3.67 \pm 0.66) \times 10^{-2}$ for $H_3 = 4.75$. The error in the exponents is the 95% confidence interval. The ensemble average is performed over 10 shifted time windows between the successive $t_w$s and 2 different trajectories, generating 20 ensembles.

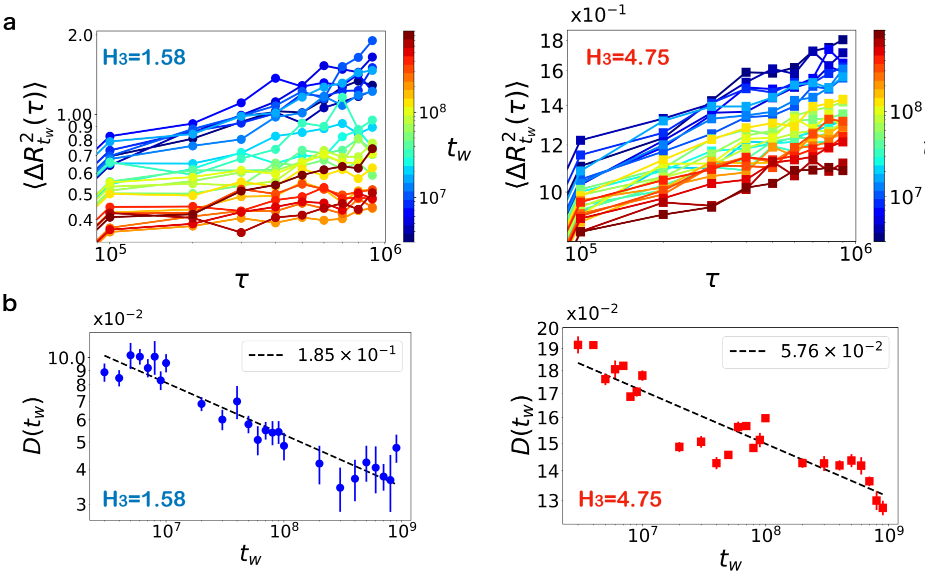


FIG. S3: **pMSD for $\epsilon_b = 4$:** (a) pMSD for low complexity sequence ($H_3 = 1.58$, left) and high complexity sequence ($H_3 = 4.75$, right). The color gradient indicates the system's age ($t_w$). (b) Same as Fig. S2, except the results are for $\epsilon_b = 4$. The dashed line is a power-law fit with exponent $\alpha = (1.85 \pm 0.28) \times 10^{-1}$ for $H_3 = 1.58$ and $\alpha = (5.76 \pm 1.09) \times 10^{-2}$ for $H_3 = 4.75$.

trap model predicts that, during aging, the system is progressively trapped in deeper energy wells over time. This is illustrated in Fig. S4a, which plots the potential energy in the simulation trajectory ($U$; see Methods section in the main text). The low complexity sequence exhibits equilibrium behavior, in contrast to the power-law increase of $-U$ observed for the high complexity sequence. Similar behavior has been found in simulations of simple glass-forming systems [S4, S5].

We also calculate the probability $P_u(t)$, representing the fraction of beads within aggregates not connected to any others (Fig. S4b with $\epsilon_b = 0$). The connection cut-off is set at $1.5\sigma$, which excludes trivial adjacent bonds in the monomers. For $H_3 = 1.58$ sequence, $P_u(t)$ demonstrates equilibration, whereas for the $H_3 = 4.75$ sequence, a power-law decay is observed as anticipated by the theoretical study [S1].

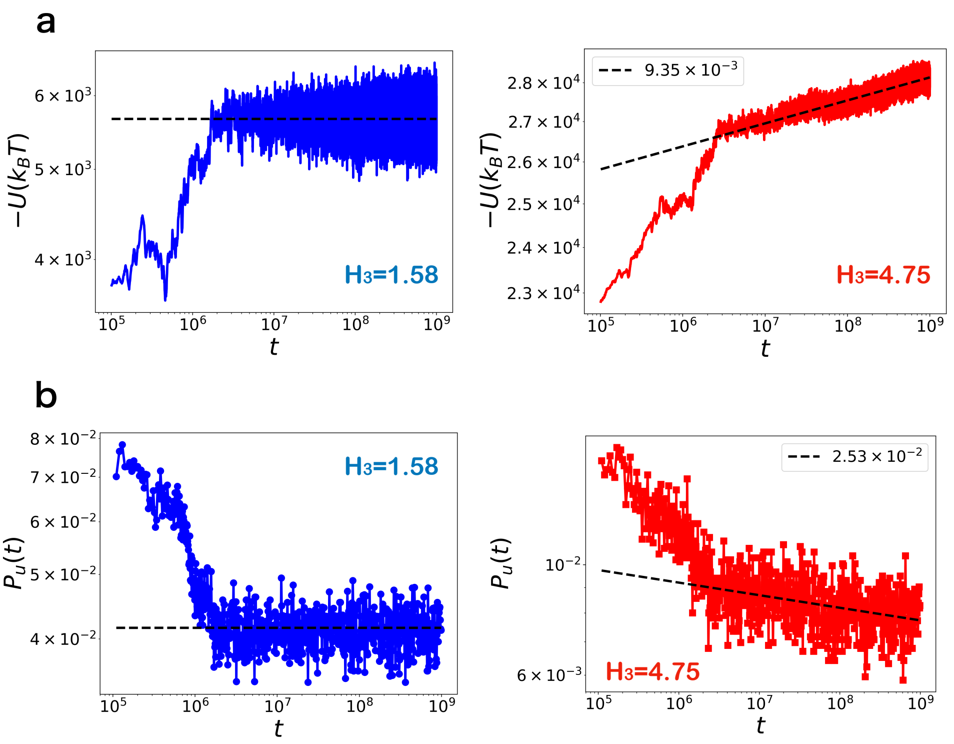


FIG. S4: **Glassy behavior in the simulation is consistent with the trap model:** (a) Potential energy ($-U$) in the simulation trajectory as a function of time for $\epsilon_b = 0$. The left and right figure show $H_3 = 1.58$ and $H_3 = 4.75$, respectively. The black dashed line is the fit for $t > 10^7$. The line is a constant for $H_3 = 1.58$ and the power-law with the exponent $(9.35 \pm 0.81) \times 10^{-3}$ for $H_3 = 4.75$. (b) The unbound probability $P_u(t)$ for $\epsilon_b = 0$. The left and right figure show $H_3 = 1.58$ and $H_3 = 4.75$, respectively. The line is a constant for $H_3 = 1.58$ and the power-law with the exponent $-(2.53 \pm 0.81) \times 10^{-2}$ for $H_3 = 4.75$. The errors of the exponents are the 95% confidence interval of the estimates.

### IV. VIDEOS

Supplementary movie 1: Initial stage ($t \leq 3 \times 10^6$ steps) of the ordered aggregate formation for the parameters $H_3 = 1.58$ and $\epsilon_b = 2$. Bead colors denote the sequence letters, with red corresponding to A, green to B, and blue to C. Initially liquid-like and disordered aggregate forms. Subsequently, an ordered structure starts to appear.

Supplementary movie 2: Initial stage ($t \leq 3 \times 10^6$ steps) of the ordered aggregate formation for the parameter $H_3 = 1.58$ and $\epsilon_b = 4$. Bead colors denote the sequence letters, with red corresponding to A, green to B, and blue to C. Ordered and elongated structure appears rapidly after aggregation.

[S1] R. Takaki, L. Jawerth, M. Popović, and F. Jülicher, *Theory of rheology and aging of protein condensates* (2023), 2303.18028.
[S2] J.-P. Bouchaud, Journal de Physique I **2**, 1705 (1992).
[S3] C. Monthus and J.-P. Bouchaud, Journal of Physics A: Mathematical and General **29**, 3847 (1996).
[S4] W. Kob and J.-L. Barrat, Physical review letters **78**, 4581 (1997).
[S5] L. Angelani, R. Di Leonardo, G. Parisi, and G. Ruocco, Physical Review Letters **87**, 055502 (2001).
